# Network nature of ligand-receptor interactions underlies disease comorbidity in the brain

**DOI:** 10.1101/2024.06.15.599140

**Authors:** Melissa Grant-Peters, Aine Fairbrother-Browne, Amy Hicks, Boyi Guo, Regina H. Reynolds, Louise Huuki-Myers, Nick Eagles, Jonathan Brenton, Sonia Garcia-Ruiz, Nicholas Wood, Sonia Gandhi, Kristen Maynard, Leonardo Collado-Torres, Mina Ryten

## Abstract

Neurodegenerative disorders have overlapping symptoms and have high comorbidity rates, but this is not reflected in overlaps of risk genes. We have investigated whether ligand-receptor interactions (LRIs) are a mechanism by which distinct genes associated with disease risk can impact overlapping outcomes. We found that LRIs are likely disrupted in neurological disease and that the ligand-receptor networks associated with neurological diseases have substantial overlaps. Specifically, 96.8% of LRIs associated with disease risk are interconnected in a single LR network. These ligands and receptors are enriched for roles in inflammatory pathways and highlight the role of glia in cross-disease risk. Disruption to this LR network due to disease-associated processes (e.g. differential transcript use, protein misfolding) is likely to contribute to disease progression and risk of comorbidity. Our findings have implications for drug development, as they highlight the potential benefits and risks of pursuing cross-disease drug targets.

## Introduction

Neurodegeneration is characterised by neuronal cell death and synaptic loss, which impacts cognition and motor function (*1*). For highly prevalent neurodegenerative disorders (NDDs), such as Alzheimer’s and Parkinson’s disease (PD), age is an important risk factor. Paired with increasing life expectancy in many countries, this means that NDDs are forecast to generate an increased social and economic burden in upcoming years (*2*). The need for therapies to halt all types of neurodegeneration has never been more pressing.

Presentation of each NDD is linked to the neuronal population affected in each disease, such as loss of dopaminergic neurons in the substantia nigra driving motor dysfunction in PD or depletion of motor neurons in the spinal cord and upper motor neurons in the cortex, which drives disability in amyotrophic lateral sclerosis (ALS) (*3*). In spite of these differences, NDDs have several overlapping features (*4–6*). These include aberrant protein homeostasis, altered energy homeostasis, DNA and RNA defects, inflammation and, ultimately, neuronal cell death (*1*). Many of these processes are interlinked. For example, aberrant protein homeostasis can cause mitochondrial stress, impacting energy homeostasis and contributing to neuronal death. Furthermore, co-pathology is the norm rather than the exception in neurodegeneration, where 90% of older adults with Alzheimer’s disease (AD) with autopsy confirmation have mixed pathology (*7*). Although aggregation of amyloid beta and tau fibrils, alpha synuclein and TDP-43 are stereotypically associated with different diseases, they frequently co-occur in the same patient, with features from as many as 7 neurodegenerative conditions reported in the same individual (*4, 8, 9*). High comorbidity rates are potentially further evidence of the overlaps in the neurodegenerative process. For example, 20-40% of patients with Parkinson’s disease (PD) go on to develop dementia (*10, 11*). Taken together, the overlapping biological pathways, and high rates of co-pathology and comorbidity suggest that while NDDs have disease-specific features, there are important overlaps in disease processes. The cause of these overlaps remains largely elusive.

Genotype-phenotype relationships of genes associated with NDDs are complex, as demonstrated by familial forms of disease. Variable expressivity is well recognised, as in the case for the pathogenic repeat expansion of *C9orf72* which can cause both amyotrophic lateral sclerosis (ALS) and frontotemporal dementia (FTD) (*12*). Furthermore, pathogenic variants in a range of genes can cause the same clinical outcome, such as dominant mutations in amyloid precursor protein, presenilin 1 and 2, all of which result in early onset AD (*13*). A further layer of complexity is added with sporadic forms of disease, which account for the majority of cases. Single nucleotide polymorphisms that increase disease risk are largely independent across neurodegenerative disorders when heritability is considered globally (*14*). Although there is evidence for local genetic correlations across diseases (*15*), these seem unlikely to be sufficient to explain the overlaps observed across NDDs.

Neuropsychiatric disorders (NPDs) also have several overlapping features, including alterations to fundamental molecular processes and symptomatology (*16, 17*). For instance, alterations to synapses and circuitry at various developmental stages have been associated with neuropsychiatric disease (*18, 19*). This potentially explains the high comorbidity rates across NPDs, with 48.6-51.0% of patients with major depression having at least one concomitant anxiety disorder and only 26.0-34.8% having no comorbid mental disorder (*20, 21*). Symptoms of NPDs also tend to co-occur: 21% of people fulfilling DSM-IV criteria for a mental disorder meet criteria for three or more other NPDs (*22*). However, unlike NDDs, NPDs have overlapping genetic architectures, sharing a large number of risk loci (*14*). The comorbidity rate and overlaps in genetics are such that NPDs were proposed to be spectral disorders (*23, 24*). Diagnostic guidelines have rather stratified diagnostic processes further, causing a still debated question of reification in NPD classification, that is, whether creation of this model for NPD classification has generated a bias in perception of NPDs (*25, 26*).

Irrespective of their origin, cross-disease targets have the potential to be highly important sites for drug targeting. To date, the strategy for identification of such targets has largely been based on shared biology. However, since it is now known that drugs with genetic evidence are over four times more likely to be approved following clinical trials (*27*), providing a genetic context is valuable. Thus, understanding how distinct risk genes can contribute to overlapping pathways and shared disease outcomes could be an essential step for cross-target identification and successful drug design. In light of this, we hypothesised that ligand-receptor interactions (LRIs) are a promising mechanism by which products of distinct genes interact to produce the same outcome. As such, disruption to either of the interactors in an LRI would result in overlapping, albeit non-identical outcomes (**Fig 1A**). Sources of disruption to LRIs could be alterations to protein conformation and affinity due to genetic variants and/or environmental factors.

**Fig 1.**
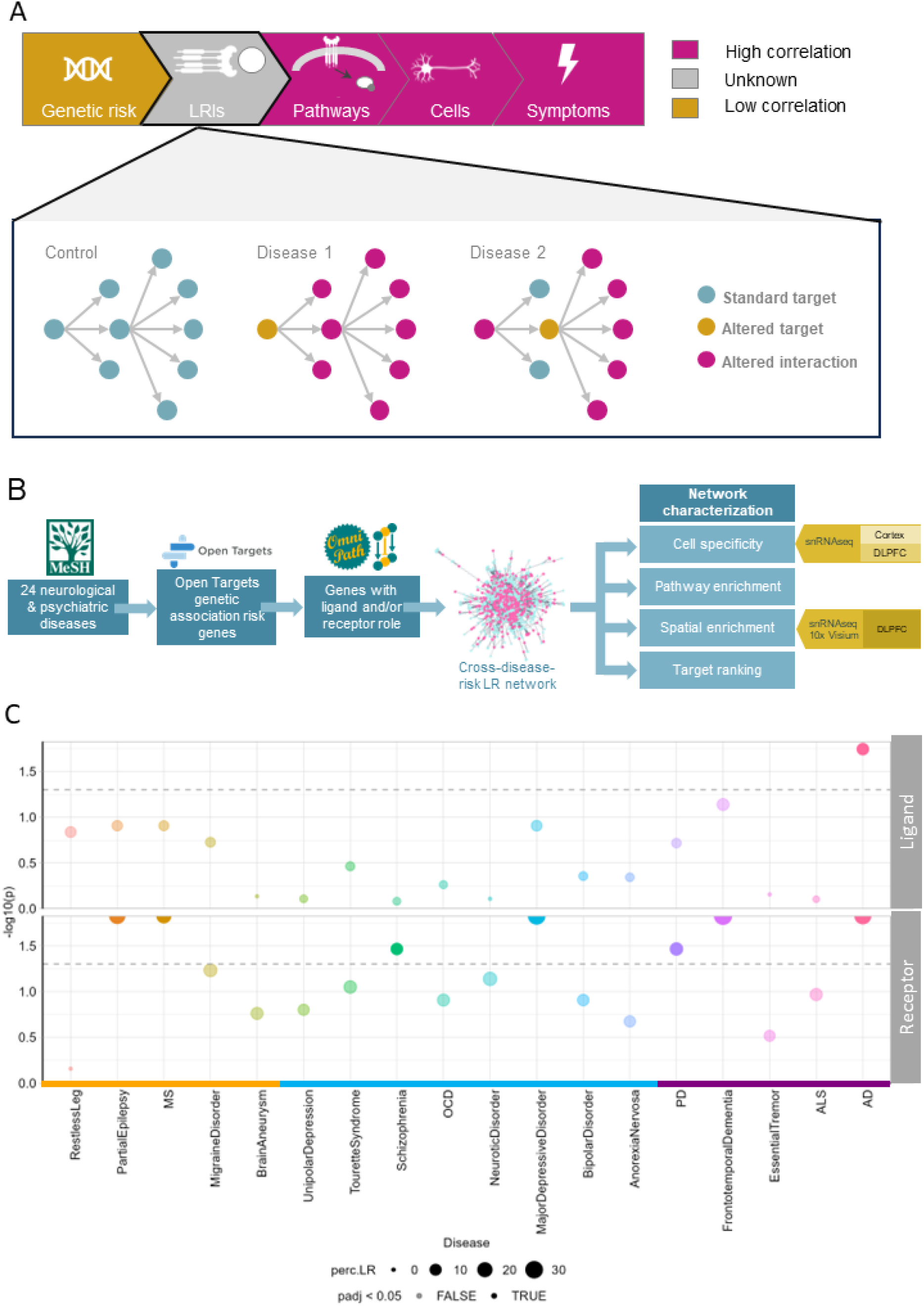
Schematic of study hypothesis, design and preliminary assessment of hypothesis. (A) Neurological and neuropsychiatric disorder have limited overlaps in genetic risk, but have abundant overlap in disease-associated pathways, cell states and symptoms. LRIs are a mechanism by which seemingly unrelated genes linked to risk of different diseases can interact to produce shared and/or overlapping outcomes. (B) Schematic illustration of study design and workflow. Briefly, neurological and neuropsychiatric diseases listed on MeSH with evidence of genetic risk associations and listed on OpenTargets were included. The genes associated with disease risk were downloaded from OpenTargets and filtered to only include genetic association scores >0.4. These genes were then filtered to only include those which have a known ligand and/or receptor role, as specified in the Omnipath database. (C) Receptors are significantly enriched among disease risk genes across 7 neurological and neuropsychiatric disorders. Ligands are significantly enriched in disease risk genes for Alzheimer’s disease.

Importantly, LRIs also tend to be highly druggable (*28*). They are often amenable to targeting by small molecules, which is essential for diseases of the central nervous system (CNS), as any treatment must be capable of traversing the blood brain barrier. Consequently, we think that characterising the role of LRIs across diseases is important. The availability of databases such as omnipathDB (*29*), which comprehensively annotates LRIs and the increasing understanding of the genetic architecture of NDDs and NPDs make this form of analysis timely. Furthermore, the availability of untargeted transcriptomics methods across cell types and tissues, can provide (1) a survey of LRIs predicted to be occurring on a cell-type level and (2) validation of LRIs that are likely to be occurring based on the location of expression (*30, 31*).

By taking into consideration the network nature of LRIs, we have found novel links between neurological diseases which are consistent with clinical observations. We also found compelling evidence that LRIs are likely to be disrupted in neurological disease. Finally, we identified and characterised a single LRI network enriched for cross-disease risk, including NDDs, NPDs and other neurological disorders, which has implications for cross-disease target design.

## Results

### Ligand-receptor interactions are expected to be disrupted in neurological disease

In order to determine whether LRIs could be a driver of co-pathology, the first hypothesis we tested was whether genes associated with neurological disease are more likely to be ligands and/or receptors (LRs) than those that are not associated with disease. We included a range of disorders including NDDs, NPDs and other neurological conditions. This resulted in the identification of 24 potential diseases of interest. For each of these diseases we identified genetically implicated gene sets using the Open Targets (https://www.opentargets.org/) resource. Only genes with genetic association risk scores > 0.4 were retained (maximum association score of 1.0) to ensure higher target confidence and only diseases with more than 10 risk genes were included. This resulted in the analysis of 18 diseases in total (**SupFig 1A-B**). We then annotated genes based on whether they are known to act as a ligand and/or receptor using the Omnipath database (*29*) (**Fig 1B**) and compared the occurrence of ligands and receptors in disease-associated genes to the occurrence of LRs amongst all protein-expressing genes expressed in the brain as detected within the Genotype-Tissue Expression (GTEx) data. Of the 18 neurological disorders tested, 7 had significant enrichment of ligands and/or receptors amongst the gene sets, namely AD, frontotemporal dementia (FTD), PD, major depressive disorder (MDD), schizophrenia (SCZ), multiple sclerosis and partial epilepsy (**Fig 1C**). The disease type with the most significant LR enrichment was NDDs. We found that 3 of the 5 tested NDDs (60%, FDR-corrected p value range 5.0×10^-2^ - 1.0×10^-4^), 2 of the 8 NPDs (25%, FDR-corrected p value range 5.0×10^-2^-1.0×10^-4^) and 2 of the 5 other neurological disorders (40%, FDR-corrected p value<1.0×10^-4^) had significant overrepresentation of LRs amongst risk genes. These results were not explained by gene set size, as exemplified by the fact that both SCZ (N = 317 genes) and FTD (N = 26 genes) had significant receptor enrichment despite having a 12- fold difference in gene set sizes. Strikingly, only AD risk genes were enriched for both ligands and receptors, while all other diseases only had significant enrichment of receptors amongst the risk genes. This overrepresentation of receptor involvement is a feature specific to neurological disorders, with disorders of other types (cardiovascular, autoimmune, cancer) either being enriched for both ligands and receptors or neither (**SupFig 1C**). This highlights the differences in genetic architecture across multiple body systems.

### Accounting for ligand-receptor interaction networks highlights disease-disease relationships

Given that receptors are enriched across risk genes for many neurological disorders, we postulated that this could drive dysfunction more broadly to cause secondary neurological disease due to their interactive nature. Specifically, disruption to the same LRIs in different diseases could generate overlaps in pathology and symptomatology. We investigated this possibility in three ways: (i) by determining whether there were significant overlaps amongst ligands and receptors themselves, (ii) by determining whether there were common LRIs affected across diseases, and (iii) by determining whether architecturally the LR networks affected in each disease were similar. We did this by comparing the real overlapping features of diseases to bootstrapped overlaps, if the real overlapping feature was in the 90th percentile of bootstrapped data, it was considered a “hit”.

Focusing on the first approach, we found that 47 out of all 153 pairwise comparisons between different diseases had direct overlaps in ligands and/or receptors with the majority of these involving NPDs (**Fig 2A**). This is consistent with the existing literature which shows that NPDs have significant overlaps in genetic risk (*14, 32*), but here we demonstrate this for LRs specifically. Out of 17 pairwise comparisons, SCZ had 12 “hits” for overlapping ligand and/or receptors, making it the disease with most “hits”. This could be expected, since SCZ has the largest number of risk-associated LRs. The disease with the second highest number of overlapping “hits” was partial epilepsy, which overlapped with all NPDs as well as several NDDs (AD, PD, FTD) and migraine disorder.

**Fig 2.**
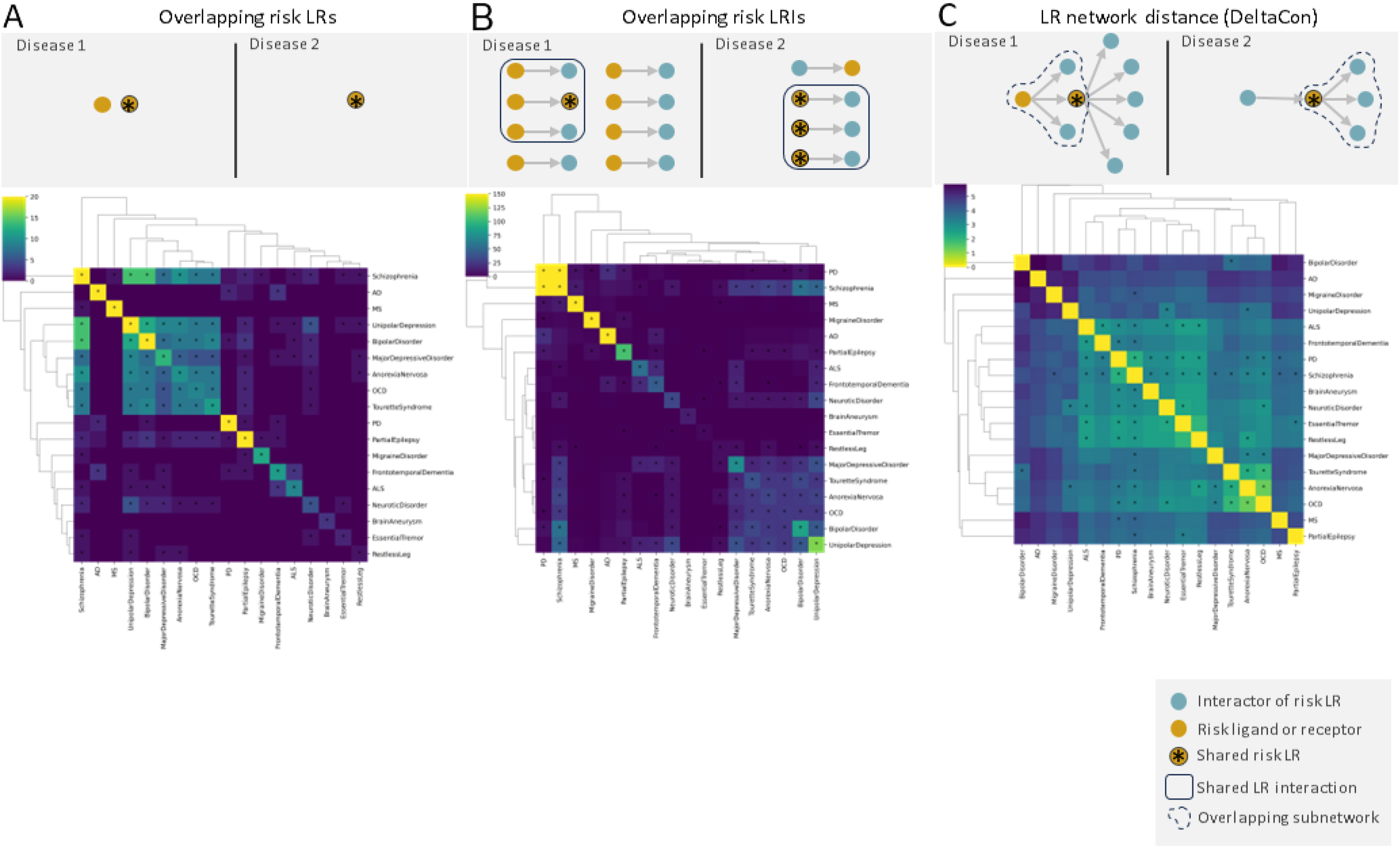
Taking into account the network nature of LRIs highlights similarities between neurological and neuropsychiatric diseases reflected in symptoms, comorbidity. (A) Ligands and receptors associated with disease risk overlap most frequently for neuropsychiatric disorders. (B) PD and schizophrenia are the disease pair with most overlaps of at-risk LRIs, i.e. interactions where at least one interactor is associated with disease risk (C) A group of 8 disorders, including neurological and neuropsychiatric conditions, has significantly similar LR networks associated with disease risk.

We investigated LR relationships further using our second approach, namely assessing how many LRIs containing risk genes each disease pair shared (**Fig 2B**). For this, an LRI was considered to contain risk genes if at least one of its interactors (ligand or receptor) was associated with disease risk. For example, in a case where a ligand was associated with one disease and an interacting receptor with another, this LRI would be considered of interest for both diseases. We found that PD and SCZ were the comparison with most LRI overlaps, with 169 overlapping LRIs. This was largely due to the gene *FYN*, which is associated with both PD and SCZ. We noted that NPDs had similar overlap profiles, with several “hits” amongst each other. Interestingly, both unipolar depression (N = 12 “hits”) and major depressive disorder (N = 9 “hits”) had the highest number of LRI “hits” with other diseases, including all NPDs, some NDDs (ALS, FTD) and other disorders (partial epilepsy, restless leg).

Finally, we assessed the similarities of LR networks associated with each diseases as a whole (**Fig 2C**). To do this we calculated the similarity of LR networks for each disease pair using the deltacon algorithm, which compares the connectivity of two networks. In this method small distances indicate a high degree of similarity between the networks and large distances indicate more distinct networks.

SCZ was the disease with the greatest LR network similarity with other diseases (N = 14 “hits”), including NPDs (OCD, Anorexia nervosa, tourette syndrome, MDD, neurotic disorder), motor disorders (Parkinson’s disease, essential tremor, restless leg syndrome, amyotrophic lateral sclerosis) and other disorders (multiple sclerosis, partial epilepsy, brain aneurysm, migraine disorder). PD was the disease with the second highest similarity (N = 10 “hits”), with disease overlaps that included movement disorders (essential tremor, restless leg syndrome), NDDs (FTD, ALS), NPDs (OCD, anorexia nervosa, neurotic disorder, major depressive disorder) and other neurological disorders (MS, partial epilepsy). Based on this analysis, we concluded that the LR networks were a promising route by which to understand relationships across multiple neurological diseases, even across disease types.

### A single ligand-receptor network connects the majority of risk-associated ligands and receptors across diseases

We found that LR networks across neurological conditions had significant network similarities in pair-wise comparisons. However, this analysis did not capture similarities across more than two diseases. Therefore, we explicitly generated and characterised the LR network architecture across all diseases within a single analysis. We foresaw three main possible outcomes: (i) each disease would have its own LR network containing large numbers of risk genes (e.g. PD-specific network), (ii) there would be an LR network for each major disease type (e.g. NDDs) with risk genes for a number of diseases included, and (iii) there would be one major LR network enriched for disease risk genes associated with multiple disease types (e.g. NDDs and NPDs). We found that the LR network we generated was highly connected and included all major disease types (namely outcome iii). Specifically, 96.3% of all disease associated LRs (based on a genetic association threshold of 0.40) were interconnected and contained within a single network (99th percentile, **Fig 3A**), which we termed the cross-disease LR network. We tested whether this network was maintained as we increased the stringency of selection of risk genes to only keep LRs with higher risk association scores. We incrementally increased the genetic association score threshold to 0.55 and 0.70, and found that the proportion of LRs in the cross-disease LR network was largely maintained. More specifically, using a genetic association score of 0.55, 94.9% of genes were still part of a major cross-disease LR network (99th percentile). Even at a very stringent genetic association score of 0.70 the majority of LRs were in the main network (65.6% of LRs, 60th percentile, **SupFig 1C**). These findings show that LRs associated with disease risk are functionally interconnected and that disruption of disease-associated LRIs in one disease would be expected to impact the broader LR network to potentially generate comorbidity.

**Fig 3.**
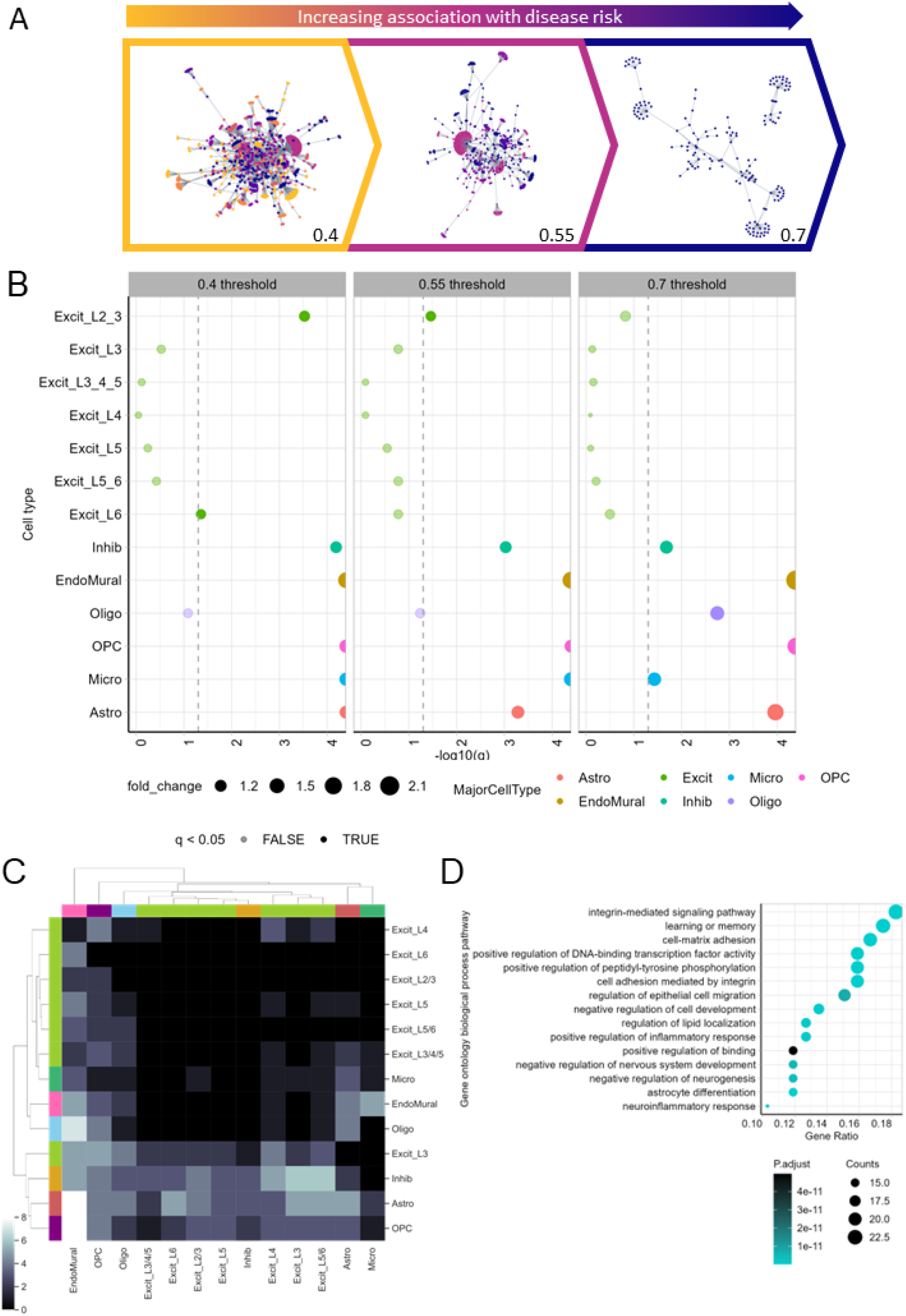
95% of LRIs associated with disease risk across 18 diseases form a single interconnected LR network expressed in glia and inhibitory neurons. (A) This major network was maintained even with increasing stringency, indicated at >0.4, >0.55 and >0.7 genetic association scores. At stringency 0.7 the network splits into two subnetworks, but the largest still has 65% of all LRIs associated with disease risk (B) Cell type enrichment analysis shows that the genes within the cross-disease risk network are enriched to glial cell types and inhibitory neurons. This becomes more significant with increasing genetic risk association score stringency (scores > 0.7) (C) The risk-associated LRIs are most frequently associated with glia-glia communication in the control dorsolateral prefrontal cortex (D) Genes in the risk-associated LR network are enriched to pathways linked to brain development, neuroinflammation.

### Glia and inhibitory neurons have a disproportionate role in cross-disease risk

Having identified a cross-disease LR network enriched for disease risk, the next step was to characterise the genes in this network (N = 865 genes). Specifically, we wanted to determine the cellular specificity of the genes and their contribution to known pathways. Focusing on the former, we used single nuclear RNA sequencing data (human cortex) and expression weighted cell type enrichment (EWCE) analysis to identify cell types of interest. We found that the risk LR network had significant enrichment for genes with high specificity for endothelial cells (FDR-corrected p value < 1.0×10^-4^), OPCs (FDR-corrected, q < 1.0×10^-4^), astrocytes (FDR-corrected p value < 1.0×10^-4^), microglia (FDR-corrected, q < 1.0×10^-4^) and inhibitory neurons (FDR-corrected p value = 2.0×10^-4^) (**Fig 3B**). An increase in stringency in the network (genetic association score > 0.70, N = 128 genes) resulted in enrichment to glia cell types alone (FDR-corrected, p value range 3.77×10^-2^ - 1.0×10^-4^). We validated and extended these findings using a second snRNAseq dataset from multiple human cortical regions that enabled finer annotation of inhibitory and excitatory neurons. Interestingly, the inhibitory neuron enrichment observed previously was replicated and shown to be specific to GABAergic VIP+ neurons (FDR-corrected, p value < 1.0×10^-4^, **SupFig 2A**), which are known to regulate behavioural circuitry (*33*). Furthermore, we observed significant enrichment of these genes to excitatory intertelencephalic neurons in layers 5/6 (FDR-corrected p value = 4.0×10^-4^).

Next, we used the ligand-receptor analysis framework (LIANA) (*34*), which infers LRIs in snRNAseq data directly, and in a data-driven manner. This orthogonal approach adds a relational dimension by predicting which cell types are likely to interact through each LR pair. Consistent with our EWCE-based findings, LIANA predicted the prominent role of inhibitory neurons and glia within the LR network. With increased stringency (genetic association score > 0.70) the main interactions were glia-glia: Astrocyte-Endomural (N = 8), OPC-Endomural (N = 8), Oligo-Endomural (N = 7), which mirrored results using EWCE (**Fig 3C**). Similarly, at a genetic association score > 0.40 we also found inhibitory neuron-inhibitory neuron interactions were frequently identified using the genetic association score of 0.40 and 0.55, which again is consistent with the EWCE analysis (**SupFig 3A-B**).

Finally, we performed pathway enrichment analysis with the genes in the risk LR network to identify biological processes of greatest interest. We found that the LR network genes had an overrepresentation of genes in pathways linked to integrin (integrin-mediated signalling pathway, FDR-adjusted p value 1.97×10^-25^; cell adhesion mediated by integrin, FDR-adjusted p value 7.84×10^-23^; cell-matrix adhesion, FDR-adjusted p value 1.51×10^-15^; positive regulation of binding, FDR-adjusted p value 4.96×10^-11^), cell development and/or neurogenesis (negative regulation of cell development, FDR-adjusted p value 6.24×10^-13^; negative regulation of nervous system development, FDR-adjusted p value 4.77×10^-12^; negative regulation of neurogenesis, FDR-adjusted p value 3.17×10^-12^; astrocyte differentiation, FDR-adjusted p value 1.21×10^-15^) and neuroinflammation (neuroinflammatory response, FDR-adjusted p value 3.20×10^-13^; positive regulation of inflammatory response, FDR-adjusted p value 3.78×10^-13^) (**Fig 3D**). Similar pathways were highlighted in the less stringent networks as well (**SupFig 3C-D**). Jointly, these findings shed light on the importance of the glial and inhibitory infrastructure of the brain in cross-disease risk.

### Risk-associated ligand-receptor interactions occur in the vicinity of excitatory neurons

LR interactions require spatial proximity of interacting partners. For membrane-bound proteins this means that the protein-expressing cells must be located close to each other for ligand/receptor binding to occur. For soluble LRs, proximity of expression makes a given interaction more likely, even if it is not essential for the expressing cells to be in contact.Therefore, we decided to characterise the spatial context of risk LR interactions. This serves as both validation that these interactions can occur in the human brain, and provide insight into the cell types which may be secondarily affected. For instance, we expect that co-localisation of neurons with pro-inflammatory interactions has a deleterious effect on neuronal survival and function. More specifically, we wanted to determine : (i) whether there are spatial cortical domains mapping to cortical layers which are more likely to be impacted by the risk LR network, and (ii) which cell types tend to co-localise near these risk interactions (**Fig 4A**).

**Fig 4.**
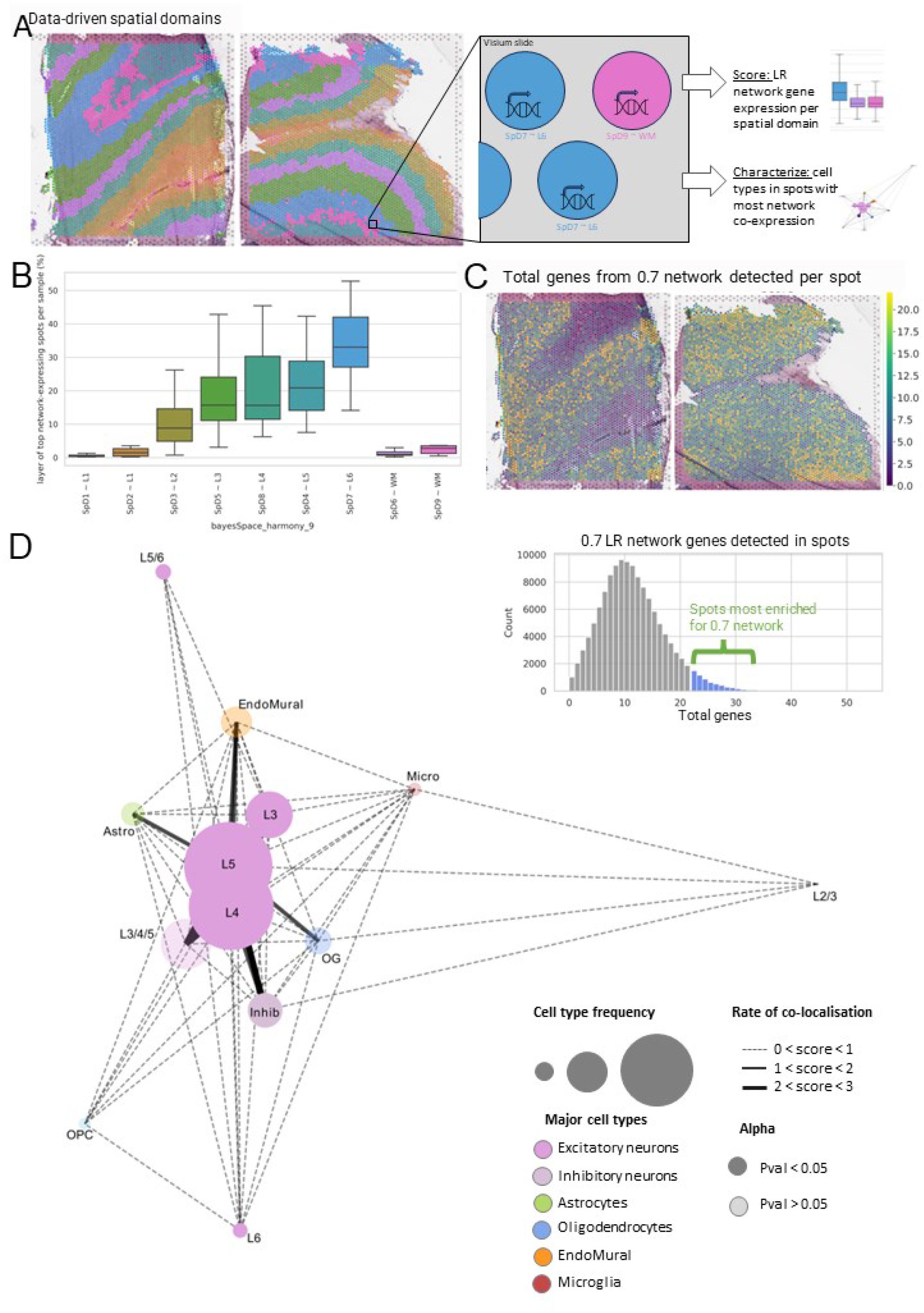
The cross-disease risk LR network is significantly enriched to layer 6 of the dorsolateral prefrontal cortex in the human brain. (A) Schematic diagram of spatial enrichment analysis. We used cortical layer-annotated dorsolateral prefrontal cortex data generated with 10x Visium to assess whether the cross-disease risk LR network was spatially enriched to any cortical layer and to assess which cell type neighbourhoods occur in the regions with most expression of the genes in the network. (B) The spots with most co-expression of the network genes (top 5%) are significantly enriched to layer 6 of the cortex. (C) The network-enriched spots are enriched for excitatory and inhibitory neurons.

To address these questions we used public spatial transcriptomic data derived from human dorsolateral prefrontal cortex and generated on the 10x Visium platform, together with paired snRNAseq DLPFC data from the same samples. Given that the Visium platform captures RNA-expression within 55 um diameter spots, we began by classifying all spots (N = 113,927) across the 30 samples available based on the expression of genes within the LR risk network. For each spot we counted the proportion of the genes within the LR network which were detected and generated a distribution of gene set detection. We focused on spots within the top 5% of the distribution and classified them as spots with high disease risk. Next, we tested whether this population of spots were more likely to be located in a specific cortical layer and found that spots were enriched within cortical layer 6. In fact, this enrichment was significant relative to all other cortical layers (Dunn’s test, 5.431×10^-58^ < p value < 1.0×10^-283^, except layer 5 (p value > 5.0×10^-2^, **Fig 4B-C**).

We also found that certain cell types were significantly enriched within spots with high disease risk. In particular, we found that inhibitory neurons and excitatory neurons from cortical layers 3-6 (for inhibitory and excitatory neurons in layer 3, 4, 5, and 6 FDR-corrected p value < 1.0×10^-3^) occurred more frequently in these spots than would be expected by chance (**Fig 4D**). This is in part consistent with the EWCE enrichment analysis, which highlighted LR enrichment to inhibitory neurons and excitatory layer 5 intertelencephalic neurons. Although we did not identify significant enrichment amongst glial cell types we noted that astrocytes and endomural cells frequently co-localised with layer 5 neurons in spots with high disease risk. Consequently, the risk LR network is most likely to impact the microenvironment of neurons in the grey matter, specifically surrounding layer 5 of the cortex.

### PD-associated genes are particularly important in the cross-disease LR network

Not all individual genes have the same potential to impact the LR network, as this is influenced by their connectivity within the network. As such, particularly in the context of drug development, identifying the most influential genes is strategic. To do this we repurposed pagerank, an algorithm used by the web search engines to rank the importance of web results, to instead rank genes within the networks. In this case, each gene was assigned importance based on (1) the number of genes it was connected to and (2) the importance of these connected genes (*35*). We then assessed which diseases these genes were associated with. Finally, we evaluated which of the highest ranking genes were part of the druggable genome (*36*) (**Fig 5A**). We noted that the five top-ranked genes, namely *RELN*, *SNCA*, *TLR4*, *APOE* and *GAL,* were part of the druggable genome (**Fig 5B**). Interestingly, 4 of these genes have been associated with PD, highlighting the importance of PD-associated genes in cross-disease risk.

**Fig 5.**
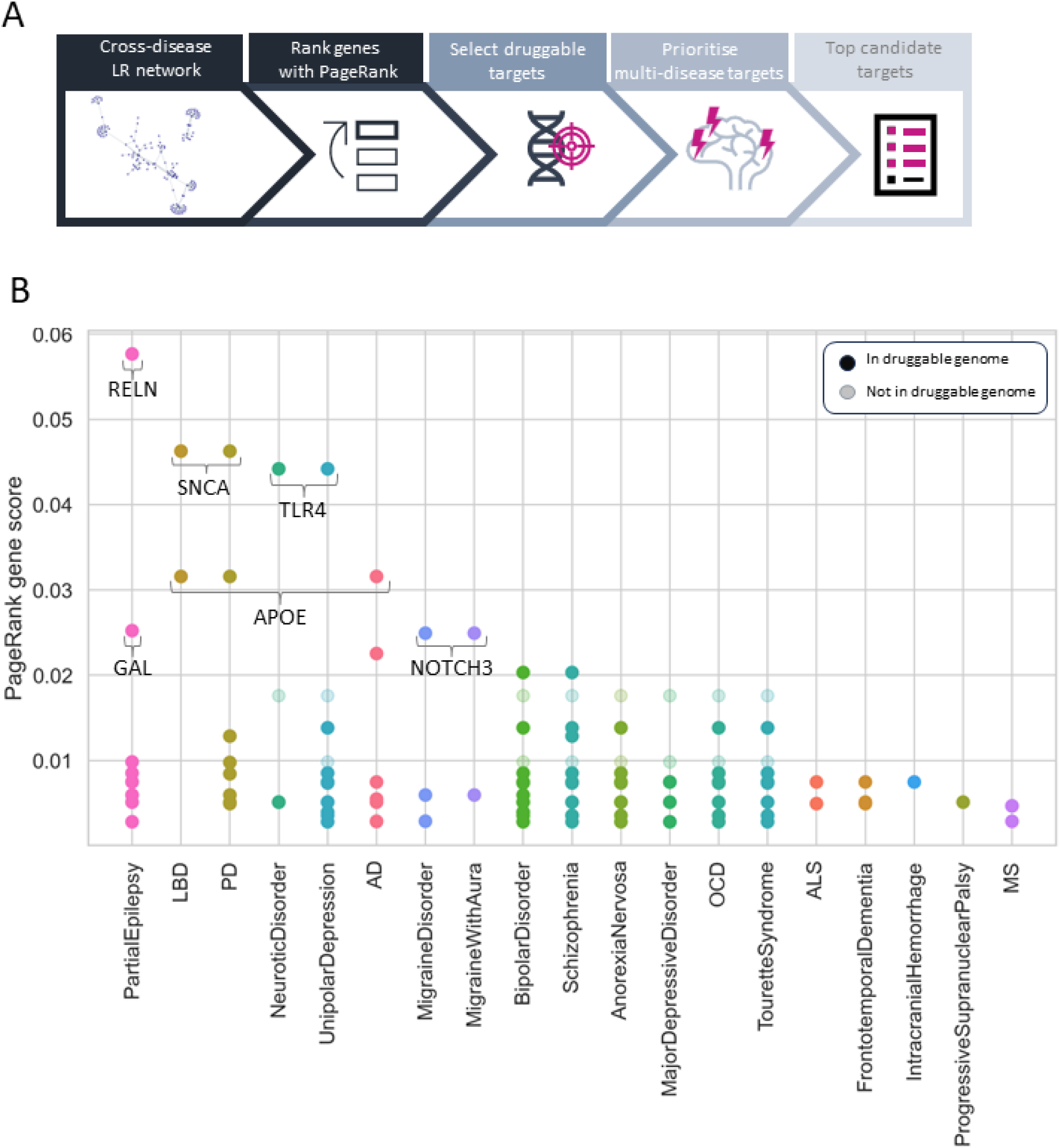
Genes associated with Parkinson’s disease are important in the cross-disease risk network and rely on accurate splicing. (A) LD-score regression analysis results for GWAS studies from AD, ALS, PD and SCZ. The cross-disease LR network (genetic association scores > 0.70) is enriched for ALS (p value < 5.0×10^-2^) and PD (p value < 1.0×10^-1^) heritability. (B) To prioritise the genes of highest importance in the cross-disease risk LR network we ranked genes based on their importance (i.e. connectivity, and the importance of connecting genes). We then assessed which of these genes are known to be druggable based on the druggable targets database and prioritised multi-disease genes. (C) The most important genes in the cross-disease LR network (genetic risk score > 0.70) have been linked to Parkinson’s disease, including SNCA (associated with PD risk), APOE (associated with PD progression), RELN and TLR4, both of which have recently been shown to be upregulated in PD. (D) We found that there is an anti-correlation between gene rank in importance and its number of transcript variants (Pearson’s R = -0.1805, p value = 1.16×10^-2^), meaning that higher ranks (closer to #1) tend to have more transcript variants. (E) The receptors associated with AD, ALS, PD and partial epilepsy have a higher number of transcript variants than would be expected (FDR-corrected, 1.32×10^-2^ < p value < 1.2×10^-3^).

## Discussion

The increasing awareness of the underlying commonalities of NDDs has fueled the pursuit of cross-disease drug targets. With several shared disease-associated pathways, frequent co-pathology and high rates of comorbidity, cross-disease targets seem like a viable and cost-effective route to tackle the lack of effective treatments for neurodegenerative disorders. In this work we explored the role of LRIs as an intermediary mechanism where the product of distinct genes interact contributing to shared outcomes in neurodegenerative and other neurological disorders.

We found that LRIs are likely to be vulnerable in several neurological diseases. Notably, when compared to diseases of other systems, genes encoding receptors were overrepresented amongst CNS risk genes. This finding may have implications for existing therapeutic efforts, which have focused on high throughput custom ligand design (*37, 38*). In fact, our findings suggest that for custom ligand design to be successful in neurological conditions, it may need to take into account disease-associated alterations in receptor conformation and affinity caused by genetic variation or alternative transcript use.

We also demonstrate that accounting for LRIs and LR networks highlights relationships between diseases which are consistent with clinical observations, but that are largely hidden when genetic data alone is considered. This extended beyond overlaps amongst neurodegenerative disorders, but demonstrated that ligands and receptor interactions are likely to contribute to overlaps between neuropsychiatric and neurodegenerative disorders. This finding is consistent with clinical data of high prevalence of neuropsychiatric comorbidities in neurodegeneration, such as depression which has a prevalence of 34-50% amongst several neurodegenerative disorders (*39–41*).

The high connectivity of ligands and/or receptors associated with disease risk highlights the degree of fine-tuning of the LRI network required for normal CNS function. We hypothesise that this network is likely to become increasingly dysregulated during the course of disease, contributing to disease progression and comorbidity. This view is supported by examples drawn from PD research. For example, *APOE,* a ligand associated with AD risk, is a key component of the LR risk network together with the receptor, *LRP1B,* which is known to bind APOE. Interestingly, genetic variation in both *APOE* and *LRP1B* have been associated with PD progression to dementia (*42*), suggesting that dysregulation of this interaction may drive additional AD-associated interactions in the network, contributing to cognitive decline and dementia in PD.

Amongst all diseases, we found that PD had particularly unique features. Although the neurodegenerative component of PD is often emphasised, our findings shed light on its neuropsychiatric component. The overlaps of PD with SCZ LRs are consistent with the known psychiatric component of PD (*43, 44*), such as psychosis (∼26-82.7% of PD patients), apathy and anhedonia (16.4-40% of PD patients) and impulse control disorders (∼14% of PD patients) (*43, 45*). Although long-term use of L-dopa may contribute to psychosis, the vulnerability of LR interactions may provide insight into why psychiatric symptoms can also occur in the prodromal phase (*46*) of PD.

We also noted that many of the genes ranked highly based on influence in the LR network have been associated with PD. This includes *FYN*, which is the highest ranking gene in cross-disease networks, and *RELN,* a secreted extracellular matrix protein associated with the migration, positioning and maintenance of neurons. Mutations to *RELN* are associated with partial epilepsy (*47*), but its anti-apoptotic role has also been of interest in the context of PD and SCZ (*48, 49*). Interestingly, *RELN* is expressed predominantly by a specialist subpopulation of inhibitory neurons in adults and is linked to the *RELN*-*DAB1* pathway which is associated with AD pathogenesis. In fact, it was recently reported to have an epistatic relationship with *APOE*, another of the high ranking targets and associated with risk of AD, PD and DLB (*50*). *SNCA* produces alpha-synuclein, a highly abundant presynaptic protein in the brain. Its misfolding and aggregation has been linked to PD and LBD, with impaired alpha-syn clearance being linked to the APOE4 variant (*51*). *TLR4* is expressed mainly by astrocytes and microglia in the brain. It has a role in neurogenesis and is linked to unipolar depression, neurotic disorder, AD, ALS, MS and PD (*52*). It is likely to be linked to inflammation through *APOE*-mediated proinflammatory signalling (*53*). These genes could be proposed as candidate cross-disease drug targets, since with minimal pharmaceutical intervention they have high potential for impact on the cross-disease risk LR network.

We note that amongst the five highest ranking genes within the LRI risk network, none currently have approved drugs. Only *TLR4* and *SNCA* have drugs currently in development. A drug targeting *TLR4*, tresatorvid, is currently in phase 3 of clinical trials (*54*), whilst two drugs targeting *SNCA*, prasinezumab and cinpanemab are in phase 2 clinical trials (*55, 56*). While the outcomes of these trials are pending, we would argue that accounting for the network nature of LR interactions as well as the directionality of LRIs associated with disease risk will be important. With high-throughput drug development pipelines and computational tools increasingly available (*37, 38*), it will become increasingly important to also leverage brain-derived cell-specific and spatial transcriptomic data . The emergence of new high resolution datasets characterising changes in gene expression , transcript use and protein isoform expression, will be especially valuable for this drug design effort.

The findings in this work must be considered in light of its limitations. Firstly, here we depend on databases, such as Open Targets and omnipathDB, being comprehensive and robust. However, there are likely to be inherent biases in the genes and LRs which are characterised and annotated. For example, in the case of OmnipathDB, other LRs may be crucial but their annotation as a ligand or receptor be unknown due to not being studied as widely.

Nevertheless, this limitation would be expected to primarily generate false negatives rather than false positives, thus not taking away the value of our findings. Secondly, we chose to use transcriptomic data for identification of (i) sites where LR interactions are likely to be occurring and (ii) to assign cell type enrichment. While transcriptomic data is currently the most sensitive approach for unbiased gene detection, we acknowledge that the correlation between gene and protein expression is poor and eagerly anticipate the wider availability of cell type-specific and spatial proteomic data.

In summary, in this study we set out to understand whether ligands and receptors were associated with cross-disease risk in neurological disorders. Indeed, we find compelling evidence that ligands and receptors are especially associated with neurological disease, and may underlie not only the overlaps in symptoms and mechanisms of neurodegenerative disorders, but also the comorbidity of neurodegenerative and neuropsychiatric disorders. By leveraging snRNA-seq and spatial transcriptomic data to better understand the cross-disease LR network, we highlight the importance of glial cells and immune processes in cross-disease risk (*57–60*). Most importantly, we believe that this work shows that a shift in CNS drug development to considering targets not only as independent genes but within their network architecture is likely to be strategic, cost effective and ultimately necessary for tackling the pressing need for treatments for neurodegenerative diseases, and beyond.

## Methods

### Systematic disease selection and genetic risk data acquisition

We selected nervous system disorders available on the Medical Subject Headings (MeSH) which in the MeSH tree structures had at most two offspring, in order to avoid generalised disease classifications. We filtered these to include only those with verifiable genetic association as demonstrated by at least 3 GWAS studies, with at least 10 genetic associations overall. This resulted in a list of 24 diseases: Alzheimer’s Disease (AD), amyotrophic lateral sclerosis (ALS), anorexia nervosa, bipolar disorder, brain aneurysm, essential tremor, frontotemporal dementia (FTD), intracranial hemorrhage, Lewy body dementia (LBD), major depressive disorder, migraine disorder, migraine with aura, multiple sclerosis (MS), narcolepsy cataplexy, narcolepsy, neurotic disorder, obsessive compulsive disorder (OCD), partial epilepsy, PD, progressive supranuclear palsy (PDP), restless leg, schizophrenia (SCZ), Tourette syndrome, unipolar depression. We then harvested from Open Targets (https://www.opentargets.org/) lists of genes with genetic association to disease risk. Because genes with a genetic score <0.1 had a small disease risk (and are therefore of less interest for drug targeting) but accounted for almost 40% of all genes, we chose to only include genes with a genetic risk score >0.1 in this analysis (**SupFig 1A**).

These gene lists were filtered to only include those with a known LR role, as annotated by the omnipath database

`(op.interactions.import_intercell_network(transmitter_params = {"categories":"ligand"}, receiver_params = {"categories": "receptor"}))`. These parameters resulted in the inclusion of both membrane bound and soluble ligands and receptors, meaning that this analysis was *not* limited to cell-cell communication networks. HLA genes and protein complexes were excluded from the analysis. Protein complexes were excluded due to the further complexity these would insert in a network analysis.

We evaluated how a change in threshold affected total LRs represented in this study, total number of diseases, the proportion of ligands and proportion of receptors represented across the data (**SupFig 1B-D**). Taking this into consideration in light of how genetic risk is scored by OpenTargets, the minimum risk score threshold we selected for downstream analysis was 0.4, as (1) we estimate that at 0.4 we are likely to capture genes with stronger evidence score in OpenTargets; (2) At this resolution we capture 19 diseases and ∼1,000 LRs; (3) At this resolution diseases with a large number of risk genes (e.g. SCZ) no longer dominate the contribution to the LRs, thus reducing bias and emphasising the cross-disease aspect of our hypothesis.

We selected 6 diseases affecting systems other than the CNS for comparison: lymphoid leukaemia, diabetes mellitus, rheumatoid arthritis, asthma, coronary heart disease and cardiovascular disease. Data was collected, preprocessed and assessed using the same methods as used for the neurological enquiry.

### Ligand-receptor occurrence in disease risk gene list

To determine whether the occurrence of ligands and receptors was enriched among disease risk genes, we used bootstrapping. For each disease, the incidence of ligands/receptors (inferred using OmnipathDB) was compared to the incidence of ligands/receptors in a list of random genes, of matching length, that were also protein-expressing genes expressed in the brain (i.e.: expression >0 in GTEx (v.7) data of brain tissues). The % of genes with ligand or receptor role of the background list was also determined using OmnipathDB LR annotation. This process was iteratively repeated (N=10,000). The p-values were calculated using

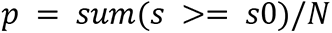

where s = bootstrapped LR incidence values, s0 = LR incidence in disease risk genes, N = number of bootstrapping iterations. P-values were corrected with false discovery rate method (fdr < 0.05) and visualised in R.

### Assessment of shared risk via ligands and receptors across diseases

We assessed the overlap in ligands and receptors linked to disease risk. For this, an adjacency matrix was constructed with counts of total overlapping ligands and/or receptors across all disease pairs. We generated a pool of genes known to act as ligands and/or receptors which did not include any disease-associated LRs (N=1,800). This was our reference pool for bootstrapping. For each disease pair comparison, we subsampled the reference LR pool to have an LR list of matching length to the real disease-implicated LRs (e.g. 300 genes for SCZ and 10 genes for PD). The overlap of these two random lists was then estimated, and the process was repeated iteratively. Because the sample pool only included 1,800 LRs and our largest disease-LR set was 300 genes, we repeated this process 12 times to allow random resampling to only have a 50% overlap in LR sampling per iteration. We then compared the bootstrapped values to the real overlapping values, reporting as of interest disease overlaps within the 90th percentile of the bootstrapped data.

### Assessment of shared LR interactions across diseases

We assessed the overlap in LR interactions affected by disease-associated genes. For this, an adjacency matrix was constructed with counts of total overlapping LRIs across all disease pairs. These values were then compared to bootstrapped data using the same method as reported for direct ligand and receptor overlap, with overlaps being reported as of interest when real values were within the 90th percentile of bootstrapped LRIs.

### Assessment of LR network similarity across diseases

For an overall analysis of the similarity between networks of different diseases (including all LR pairs where at least one interactor is linked with that disease), we calculated network similarity with the netrd python library implementation of DeltaCon (*61*). With DeltaCon, the similarity of two networks is given in the form of a distance score, where closer networks (lower score) are more similar and more distant networks (higher score) are more distinct. The algorithm assumes that both networks have the same nodes but evaluates similarities in edges. Therefore, all diseases had the same genes but only relevant edges associated with disease risk were logged and compared. The final disease-associated networks for each disease were systematically compared in a pair-wise comparison, with scores visualised in a symmetrical clustermap using seaborn. The likelihood of random occurrence of network closeness for each disease pair was estimated using the bootstrapping method used for direct LR overlap and LRI overlap, with distances in the top 90th percentile being considered of interest.

### Cross-disease risk network analysis

Using the list of all genes associated with disease risk with a known LR role, all LR interactions listed in OmnipathDB where at least one interactor was associated with neurological disease were fetched. This was logged as an adjacency matrix and visualised as a network using networkx. This was repeated longitudinally across risk scores (genetic risk score > 0.4, >0.55 and >0.7 in Open Targets). At higher risk thresholds there is a breakdown of the network structure, and therefore these analyses were not extended further.

### Major core network analysis

Community detection was performed using the louvain algorithm and the largest interconnected network was determined based on the community with the largest number of genes. Network visualisation was used to ensure the leiden parameters were consistent with the selection of interconnected networks and not further subdivision of the network.

In order to determine whether the size of the largest network was statistically significant, the size of the network was compared to bootstrapped simulated data. For this, a random list of matching length of the major network for each resolution was iteratively generated (n=10,000) with genes expressed in the brain (as determined by GTEx) and which have a LR role (as determined by OmnipathDB). This was performed for risk genes of increasing stringency (genetic association > 0.4-0.7). The size of a network was considered significant if it was in the bottom 5th percentile or top 95th percentile of the bootstrapped distribution.

### Cell type gene expression enrichment analysis

Cell type enrichment of the cross-disease network was performed similarly to previous reports (*62, 63*). We used two separate datasets for estimation of cell type enrichment: the Human Multiple Cortical Areas SMART-seq dataset (freely available through the Allen Brain Atlas data portal, https://portal.brain-map.org/atlases-and-data/rnaseq)and a dorsolateral prefrontal cortex snRNAseq dataset (*64*) (now publicly available), as references.

The Allen brain dataset includes single-nucleus transcriptomes from 49,495 nuclei across multiple human cortical areas, including the middle temporal gyrus, anterior cingulate cortex, primary visual cortex, primary motor cortex, primary somatosensory cortex, primary auditory cortex. Cell type annotations were generated at two resolutions using the Allen Brain-provided cell subclasses. A total of 1,985 nuclei were labelled as “outlier calls” and were removed during generation of the celltype dataset. We used the function fix_bad_hgnc_symbols() (R package EWCE, version 0.99.3, https://bioconductor.org/packages/release/bioc/html/EWCE.html) to remove any symbols from the gene-cell matrix that were not official HGNC symbols. A total of 30,792 genes were retained. We then used the function drop_uninformative_genes() (R package EWCE, version 0.99.3, https://bioconductor.org/packages/release/bioc/html/EWCE.html), which removes “uninformatic genes” to reduce compute time in subsequent steps. The following steps were performed:

- Drop non-expressed genes (n=1,263). This step removed the genes that are not expressed across any cell types
- Drop non-differentially expressed genes (n=6,304), which removes genes that are not significantly differentially expressed across level 2 cell types with an adjusted p-value threshold of 1e-05.

Finally, we used the function generate_celltype_data() from the R package EWCE (version 0.99.3, https://bioconductor.org/packages/release/bioc/html/EWCE.html) to generate the celltype dataset. This dataset can be accessed at: https://github.com/RHReynolds/MarkerGenes (version 0.99.1, DOI: 10.5281/zenodo.6418604).

The DLPFC dataset includes single-nucleus transcriptomes from 54,394 nuclei from the human DLPFC without neurological disorder. Preprocessing and annotation of this dataset is reported elsewhere (*64*). We used the function generate_celltype_data() from the R package EWCE to generate the celltype dataset.

For both datasets cell-type enrichment was calculated using Expression Weighted Cell Type Enrichment (EWCE)(*65*). The goal of this analysis was to determine whether the genes of interest had significantly higher expression in certain cell types than might be expected by chance. Bootstrap gene lists controlled for transcript length and GC-content were generated with EWCE iteratively (n=10,000) using “bootstrap_enrichment_test()” function. This function takes the inquiry gene list and a single cell type transcriptome data set and determines the probability of enrichment of this list in a given cell type when compared to the gene expression of bootstrapped gene lists; the probability of enrichment and fold-change of enrichment are the returned. P-values were corrected for multiple testing using the fdr method.

### Gene Ontology enrichment analysis

Cluster Profiler was used to calculate overrepresentation of pathways associated with the genes in the cross-disease LR network across thresholds in R 4.0.2. We obtained the entrez ID for the genes in the cross-disease networks using the library org.Hs.eg.db. When running the Gene ontology enrichment analysis, we selected the ‘Biological Process’ ontology with minimum GSS size = 50 and max GSS size = 300 and a p and q value cutoff of 0.01 (values corrected with Bejamini-Hochberg method). The pathways were ranked from the lowest adjusted p value to the highest, and the top 15 pathways were visualised using ggplot2.

### Cell-cell interaction prediction in single nuclear RNA sequencing data with LIANA

Liana was used to predict LR interactions in the DLPFC (*34, 64*). We used these results to verify which cell types were most likely to be communicating via cell-cell interactions in this dataset. This was performed for the risk network at risk score > 0.40, >0.55 and >0.70. The results were summarised in a heatmap generated with the python package seaborn.

### Spatial domain enrichment analysis

We used spatial transcriptomic data from the dorsolateral prefrontal cortex (*64*) generated with 10x Visium technology to assess whether the cross-disease LR network was enriched to any region of the cortex. We chose to perform this using the annotation recommended by the authors bayespace harmony-corrected sp09. These are data-driven clusters which largely correspond to conventional cortical annotation, except for the division of WM into two distinct spatial domains and layer 2 of the cortex into two spatial domains.

Due to the 10x Visium technology, the smallest mappable spatial unit of a given section is a spot with 55 um in diameter. For every spot covered with brain tissue we assessed how many genes from the risk network were detected. We summed the total number of genes in the risk network for all spots and selected the spots with the top 2% genes detected as the spots with most likelihood of expression of the risk network. We then assessed which spatial domain (i.e. cortical layer) these spots were located. We found that layer 6 had the highest mean number of risk spots. We compared the distributions using the Kruskal-Wallis test, followed by statistical comparison using Dunn’s test in R 4.0.2.

### Spatial neighborhood characterisation

To profile the cellular neighborhood of regions enriched for the cross-disease LR network, similarly to the spatial domain enrichment analysis, we first filtered all spatial transcriptomic spots to only include those in the top 98th percentile with the highest number of co-expressed genes from the network. We then used cell type deconvolution results generated with cell2location (see Huuki-Myers et al, 2023 for deconvolution details) to determine the top 3 cell types most likely to be in these spots. We constructed an adjacency matrix to record co-localisation occurrences for each cell type pair. The matrix was then normalised such that, when converted to a network, the sum of the edges connecting to each cell type is 3 (e.g. cell type pair that always co-localised would have a connecting edge of 3). This data was then used to construct a cellular co-localisation network, which shows which cell types are most frequently present in the spots enriched for the network of interest, as well as which cell types tend to co-localise.

### Gene importance ranking with PageRank

Genes were ranked by importance in the network using the networkx implementation of the PageRank algorithm. This algorithm is used by the web search engine to evaluate importance of results but is transferable to network analysis. It ranks the nodes (in this case, genes) and assigns a weight to their importance in the network based on (1) the number of nodes it is connected to and (2) the importance of the nodes it is connected to (*35*). This analysis was run using default parameters recommended by networkx, and was performed for the risk network at three stringency levels (risk network at risk score > 0.40, >0.55 and >0.70).

### Assessment of network overlap with the druggable genome

In order to assess druggability of the cross-disease LR network we used the available druggable genome annotations, including inclusion in the druggable genome and whether the gene is targetable by a small molecule (*36*). These annotations were used for visualisation of PageRank results.

### Data availability

All databases and data sets used in this work are publicly available. Databases:

- Disease tree - Medical Subject Headings (MeSH, https://www.nlm.nih.gov/mesh/)
- Genetic association scores - OpenTargets (https://www.opentargets.org/)
- Ligand-Receptor role annotation - OmnipathDB (https://omnipathdb.org/)
- Brain-expressed gene list reference for bootstrapping - Genotype-Tissue expression (GTEx) data, v8 RNAseqQC v 1.1.9 gene median (https://gtexportal.org/home/)

Datasets:

- Single nucleus RNA sequencing data multiple cortical regions (EWCE) - Allen Brain Atlas Human Multiple Cortical Regions SmartSeq (https://portal.brain-map.org/atlases-and-data/rnaseq/human-multiple-cortical-areas-smart-seq), data in EWCE-compatible format available at https://github.com/RHReynolds/MarkerGenes
- Single nucleus RNA sequencing and spatial transcriptomics data for the DLPFC are available through the SpatialLIBD package (*66*)
- Single nucleus RNA sequencing dorsolateral prefrontal cortex LIANA analysis - https://github.com/LieberInstitute/spatialDLPFC

### Code availability

All custom code used for this work is publicly available at https://github.com/mgrantpeters/LR_project [repository to be made public upon publication].

## Acknowledgments

MGP, AFB, JB, NW, SG, and MR were funded by Aligning Science Across Parkinson’s [Grant numbers: ASAP-000478 and ASAP-000509] through the Michael J. Fox Foundation for Parkinson’s Research (MJFF). For the purpose of open access, the author has applied a CC-BY public copyright license to all Author Accepted Manuscripts arising from this submission. RHR is currently employed by CoSyne Therapeutics (Lead Computational Biologist). All work performed for this publication was performed in her own time, and not as a part of her duties as an employee.

## Contributions

Conceptualization: MGP, MR

Methodology: MGP, MR, SGR, JB

Software: MGP, AH, RHR, AFB

Validation: MGP, AFB

Formal Analysis: MGP, AH

Investigation: MGP, MR

Resources: MGP, MR

Data Curation: MGP, LHM, NE, LCT, KM, RHR

Writing-original draft: MGP, MR, JB

Writing-review and editing: MGP, MR

Visualization: MGP, BG

Supervision: MR

Project administration: MGP, MR

Funding Acquisition: MR, NW, SG

