## Supplemental figures for "Network nature of ligand-receptor interactions underlies disease comorbidity in the brain"

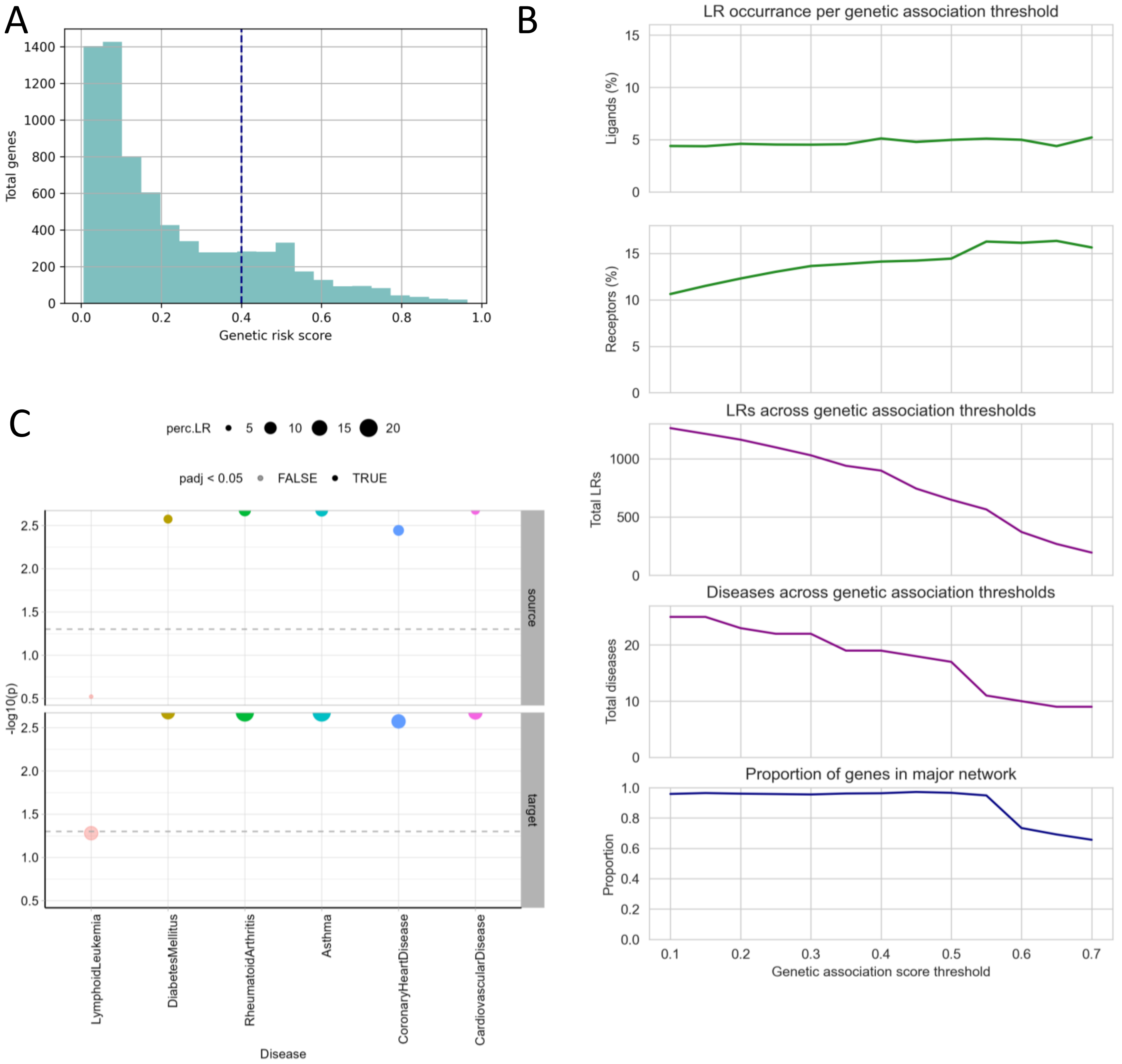


**Supplementary Figure 1:** Selection of gene risk threshold. (A) Histogram of distribution of genetic risk scores for diseases included in this study. (B) Characterisation metrics across genetic risk score thresholds. (C) ligand-receptor incidence for diseases affecting systems other than the central nervous system.

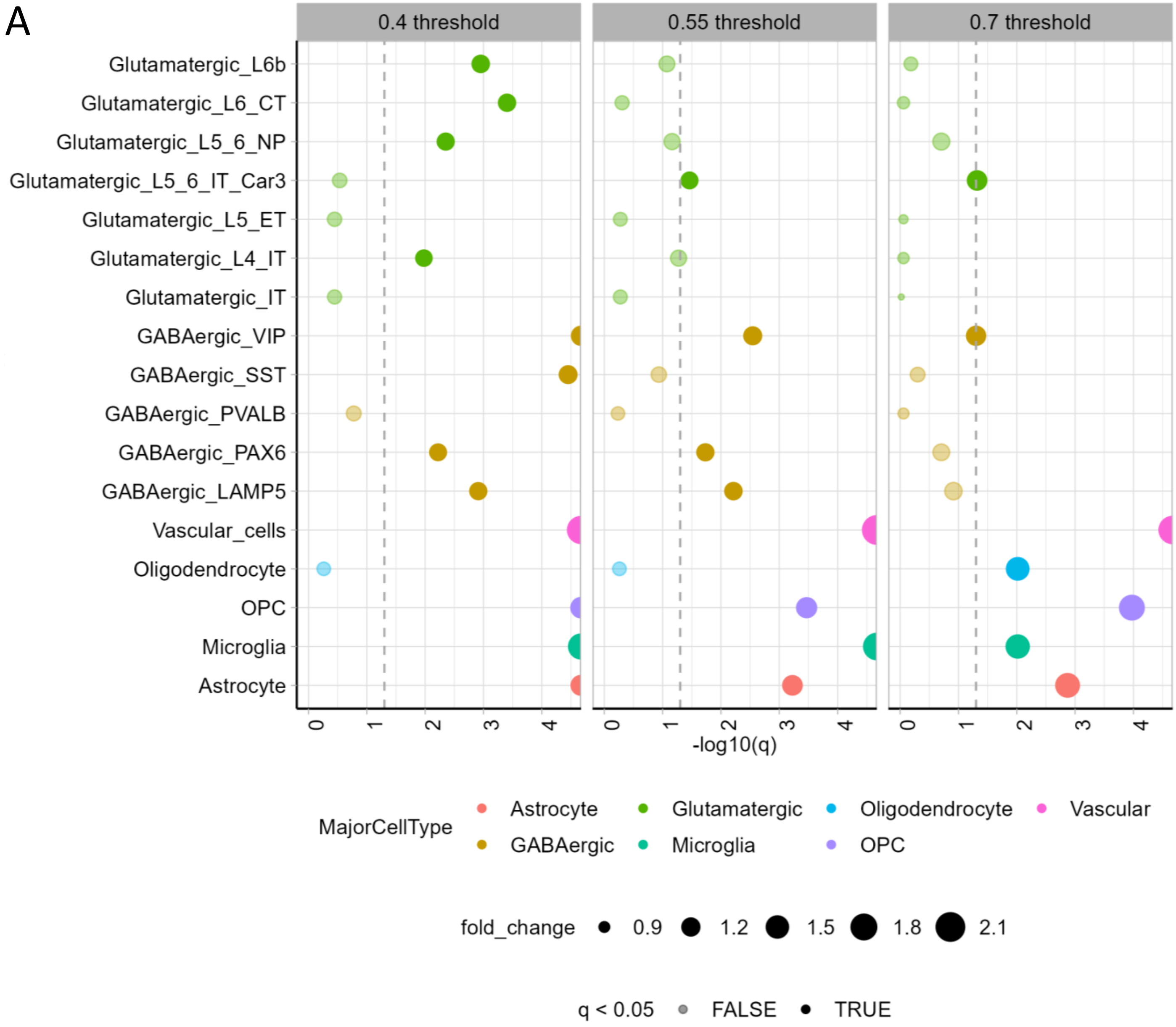


**Supplementary Figure 2:** EWCE of cross-disease LR risk network in Allen Brain Atlas multiple cortical regions data.

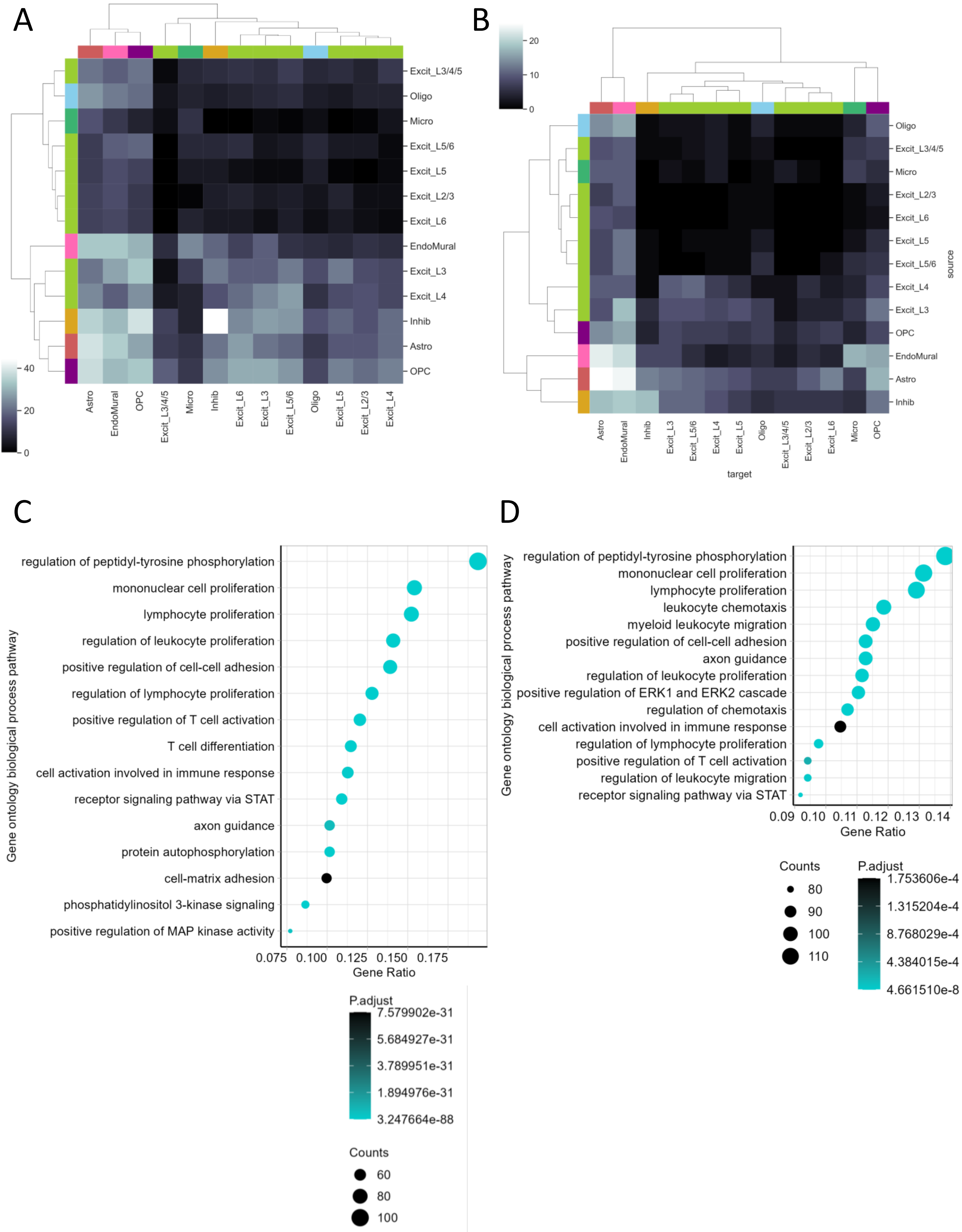


**Supplementary Figure 3:** Characterisation of LR interactions across gene risk thresholds (A) LIANA data-driven LRI inference for LRs in 0.4 threshold cross-disease LR risk network (B) LIANA data-driven LRI inference for LRs in 0.55 threshold cross-disease LR risk network (C) Biological process gene ontology pathway enrichment in 0.4 threshold cross-disease LR risk network (D) Biological process gene ontology pathway enrichment in 0.55 threshold cross-disease LR risk network

A

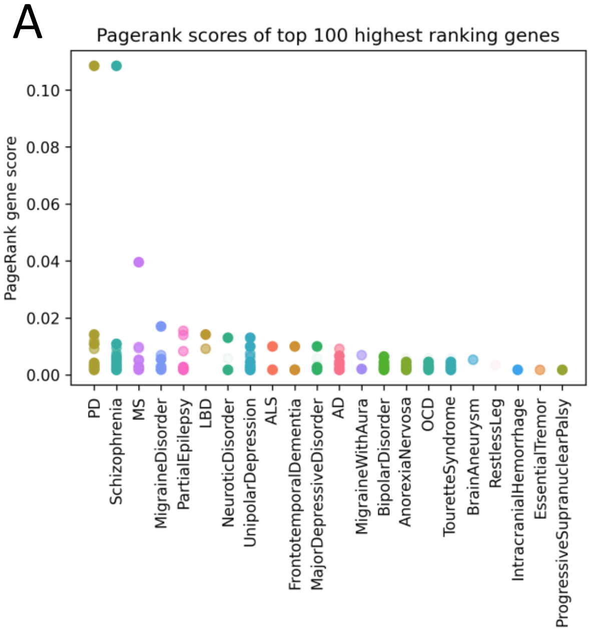


**Supplementary Figure 4:** Pagerank analysis ranking genes based on their importance (A) Pagerank results in 0.55 threshold cross-disease risk LR network
